# Insights into B Cell and Antibody Kinetics Against SARS-CoV-2 Variants Using Mathematical Modelling

**DOI:** 10.1101/2023.11.10.566587

**Authors:** Suzan Farhang-Sardroodi, Xiaoyan Deng, Stéphanie Portet, Julien Arino, Morgan Craig

## Abstract

B cells and antibodies are crucial in protecting against infections like SARS-CoV-2. However, antibody levels decline after infection or vaccination, reducing defences against future SARS-CoV-2 infections. To understand antibody production and decline, we developed a mathematical model that predicts germinal center B cell, long-lived plasma cell, memory B cell, and antibody dynamics. Our focus was on B cell activation and antibody generation following both primary and secondary SARS-CoV-2 infections. Aligning our model with clinical data, we adjusted antibody production rates for germinal center B cells and plasma B cells during primary and secondary infections. We also assessed antibody neutralization against Delta and Omicron variants post-primary and secondary exposure. Our findings showed reduced neutralization against Omicron due to its immune evasion. In primary and secondary exposures to Delta and Omicron, our predictions indicated enhanced antibody neutralization in the secondary response within a year of the primary response. We also explored waning immunity, demonstrating how B cell kinetics affect viral neutralization post-primary infection. This study enhances our understanding of humoral immunity to SARS-CoV-2 and can predict antibody dynamics post-infection or vaccination.

## Introduction

B lymphocytes, called B cells, are integral components of the adaptive immune system and contribute significantly to the human body’s defence mechanisms. These specialized cells are central to the immune response, particularly in their role as antibody producers [1]. These antibodies, also known as immunoglobulins, patrol the bloodstream and tissues, acting as a frontline defence by specifically binding to foreign pathogens and inhibiting the harmful effects of these invaders [2]. Antiviral antibodies include two distinct categories, neutralizing and non-neutralizing antibodies, each governed by unique mechanisms of action. Through particular binding, neutralizing antibodies (Nabs) have the power to entirely prevent the virus from entering host cells, halting viral particles in their tracks and effectively acting as a robust shield against infections [3–5].

The study of antibody interactions and the dynamics of B cell responses is complex. Mathematical modelling allows for insights into these intricacies, that are complementary to experimental and clinical research. Mathematical models have a well-established track record of being employed in various domains, including biology [6–9], medicine [10–12], and oncology [13–17]. They offer a simplified quantitative and predictive framework for understanding complex systems, enabling the exploration of causal relationships and mechanistic insights. Such models bridge the molecular intricacies of B cell responses and their implications in the broader context of host-pathogen interactions, allowing us to decipher the underlying principles governing immune responses. In the study discussed in [18], an initial mathematical model was proposed that integrated B cells. This model encompassed four specific types of B cells: target cells, proliferating cells, plasma cells, and memory cells. Subsequent studies expanded this model, incorporating normal B cells, memory B cells, and long-lived plasma cells to investigate the contributions of memory B cells to the secondary immune response [19]. Other studies have explored how the immune response depends on the dynamic activation of lymphocytic agents, such as T and B cells, and the interplay of signalling molecules like interleukin-2 (IL-2) and interleukin-4 (IL-4) [20]. In the research conducted by Keersmaekers et al. [21], a novel approach was employed by integrating ordinary differential equation (ODE) models with mixed effects models to examine longitudinal vaccine immunogenicity data. Utilizing B-cell and T-cell datasets from a herpes zoster vaccine study, the authors introduced ODE-based mixed-effects models, providing a valuable framework for vaccine immunogenicity data analysis and the evaluation of immunological differences between various vaccines. A comprehensive model combining the humoral immune response and the germinal center (GC) reaction has also been developed, capturing critical processes involved in immunoglobulin-G (IgG) production [22].

In this study, we used mathematical modelling to analyze the dynamic processes of B cell activation, antibody generation, and their intricate interplay with viral pathogens. Beginning from the established viral dynamics model delineating the virus-host interaction, coupled with the innate immune response, initially introduced by [23], we expanded this model by incorporating the neutralization effects of antibodies against viral particles and describing the proliferation of B lymphocytes, their differentiation into plasma and memory B cells, and the subsequent generation of antibodies following primary and secondary infections. Our primary focus centred on investigating SARS-CoV-2 variants, notably Delta and Omicron; however, it is worth noting that our model can simulate B cell activations in response to various viral infections. Our overarching goal was to construct a model capable of faithfully simulating the humoral response, guided by clinical findings. By comparing our model’s antibody predictions with clinical data from hospitalized patients, we refined our estimations of antibody generation rates by germinal center and plasma B cells. We performed a global sensitivity analysis to discern the humoral responsiveness to model parameters in primary and secondary immune responses by computing Spearman’s rank correlation coefficient between peak antibody concentrations and model parameters. The outcome of this analysis notably highlights the pronounced sensitivity of the primary antibody response to the antibody generation rate by germinal center B cells. In contrast, the secondary antibody response in its contribution shows equal sensitivity to both germinal center B cells and plasma B cells. A noteworthy insight from our findings is the need for elevated antibody generation rates to achieve equivalent antibody levels in the secondary response as observed in the primary response. We then explored antibody neutralization effects against the Delta and Omicron variants of SARS-CoV-2 within both primary and secondary immune responses. While Omicron and Delta both elicited comparable antibody levels, the former was associated with higher viral load levels and diminished neutralization efficacy. Lastly, we explored the consequences of reduced neutralization (either through viral-specific immune evasive properties or due to waning antibody concentrations) by studying re-exposure scenarios at varying intervals post-primary infection.

## Methods

In this study, we introduced mathematical models to analyze the immune response to SARS-CoV-2 infection. We begin by introducing a model (referred to as Model One, Eqs. (1)) focusing on unravelling the intricate dynamics of viral replication and the innate immune responses that are triggered upon infection. This model is primarily based on the work of [23]. Subsequently, we delve into the dynamics of the primary humoral response in our second model (Model Two, Eqs. (2)), where we present a novel mathematical framework to elucidate this essential aspect of the immune response.

We further investigate re-exposure and its impact on the secondary immune response in our newly developed third model (Model Three, Eqs. (3)). Finally, we enhance our comprehension of viral load dynamics and immune response interactions by introducing an additional neutralization function for viral load dynamics into Model One (Eq. (4)). Together, these models provide valuable insights into critical aspects of infection and immunity, offering a comprehensive exploration of the immune response to SARS-CoV-2.

### Mathematical model of viral dynamics and innate immune response

We used a simplified version of the model presented in [23] to predict SARS-CoV-2 infection within a host. This model considers a population of susceptible lung cells (*S*(*t*)) that can be infected (*I*(*t*)) by SARS-CoV-2 viral particles (*V* (*t*)). When infected, cells secrete unbound type I interferon (*F*_*u*_(*t*)), which reduces viral infection and makes the cells refractory to the virus (*R*(*t*)). Infected cells can either undergo virus-induced lysis or be eliminated by the immune response, leading to cell damage or death (*D*(*t*)). The entire model is represented by Eqs. (1).

#### Model One

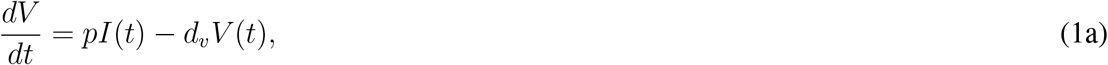

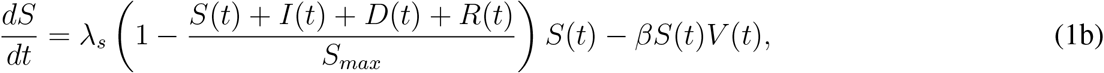

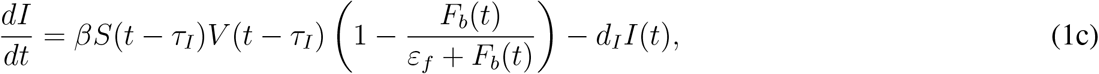

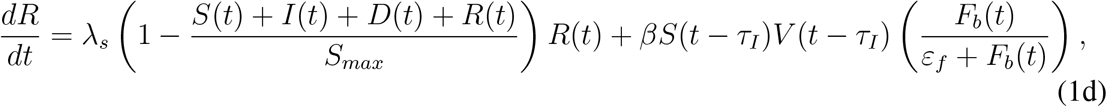

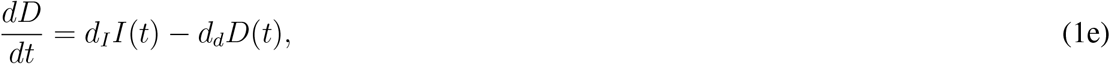

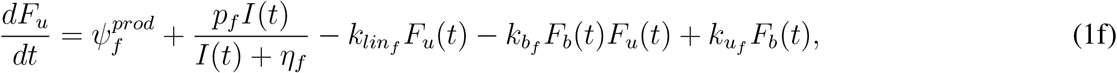

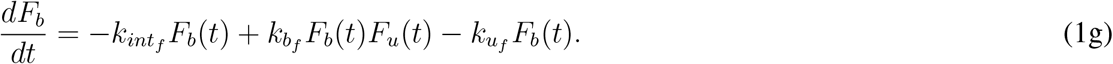

In this submodel, viral particles (*V* (*t*)) are produced by infected cells at rate *p* and are cleared through degradation at per capita rate *d*_*v*_, which accounts for all contributions to viral clearance. Susceptible epithelial cells (*S*(*t*)) proliferate logistically with a per capita proliferation rate *λ*_*s*_ to a carrying capacity of *S*_*max*_. These cells become infected (*I*(*t*)) at a rate *βV* (*t*). Resistant cells (*R*(*t*)) proliferate at rate *λ*_*s*_, which is equal to that of susceptible cells. The concentration of interferon (IFN) determines the number of cells that become refractory to infection and the number that become productively infected, controlled by the half-effect parameter *ε*_*f*_ [23]. Following an eclipse phase lasting *τ*_*I*_ hours, productively infected cells (*I*(*t*)) produce virus particles and undergo virus-mediated lysis at a rate *d*_*I*_. Dead cells (*D*(*t*)) accumulate through infected cell lysis *d*_*I*_ and disintegrate at a rate *d*_*d*_, as observed in rapid cell death [24].

The Michaelis-Menten expression 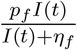 in Eq. (1f) represents the production of unbound interferon (*F*_*u*_) by infected cells in response to the infection of target cells, *I*(*t*). The parameter *p*_*f*_ characterizes the maximum rate at which unbound interferon is generated by infected cells. This maximum rate occurs when the concentration of infected cells is significantly higher than *η*_*f*_. In contrast, the parameter *η*_*f*_, known as the half-effect concentration, defines the point at which the interferon production rate reaches half of its maximum value. It plays a crucial role in determining how sensitively the production rate responds to fluctuations in the concentration of infected cells. Parameters 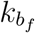 and 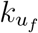 represent the binding and unbinding rates of IFN-I, respectively, while 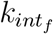 and 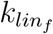 are the internalization and elimination rates of bound cytokine. Finally, the parameter 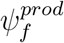 accounts for the production of IFN by macrophages and monocytes, which are not explicitly modelled in this system; for more details, refer to [23]. Table 1 provides a comprehensive list of all model parameter values along with model variables and their corresponding initial values.

**Table 1:**
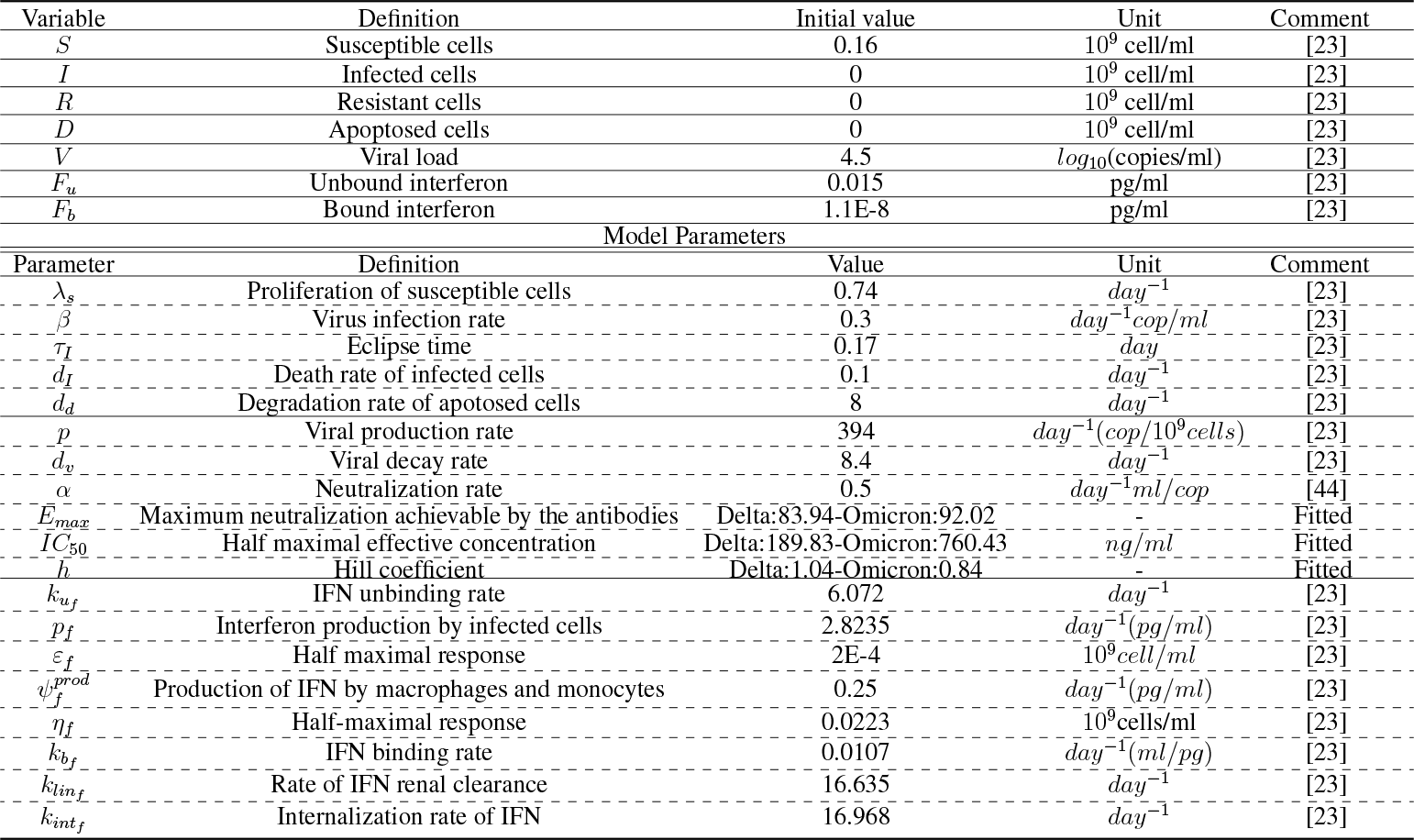
Initial Values and Parameter Settings for Innate Immune Response to SARS-CoV-2 Infection (Eqs. (1) and (4))).

### Mathematical model of the primary humoral response

Humoral immunity is a specific immune response characterized by the production of antibodies by B lymphocytes. When B cells bind to infectious agents through their surface receptors (BCRs), they release antibodies, which either neutralize or exhibit non-neutralizing effects on the antigen. In this study, we developed a mathematical model to describe the adaptive immune response, focusing on B cell and antibody-mediated immunity. Upon encountering an antigen on follicular dendritic cells in secondary lymphoid organs, naïve B cells present the antigen to T cells at the T cell-B cell border. This interaction leads to the activation, proliferation, and differentiation of naïve B cells into germinal center B cells. B cell maturation can occur within germinal centers (GCs) where activated B cells integrate immune signals, including cytokines like interleukin-4 released by follicular T cells. This process gives rise to long-lived plasma and memory B cells, which provide protective immunity by circulating in the blood or migrating to effector sites. IL-4 plays a crucial role in GC B cells’ maturation and self-renewal processes, and its absence hinders the proper formation and self-renewal of GC B cells [25]. Memory B cells and long-lived plasma cells are responsible for lifelong B cell-mediated protection against diseases [26]. Our working assumption regarding the primary response is based on a system that begins without a triggered immune response.

Based on the mechanisms described above, we developed a mathematical model of antibody production, as illustrated in Fig. 1. We explicitly considered activated B cells (*B*(*t*)), GC B cells (*B*_*g*_(*t*)), plasma B cells (*P* (*t*)), memory B cells (*M* (*t*)), neutralizing antibodies (*A*(*t*)), T follicular helper cells (*T* (*t*)), and the central cytokine interleukin-4 (*Il*(*t*)). The model consists of the following system of seven nonlinear delay differential equations:

**Figure 1.**
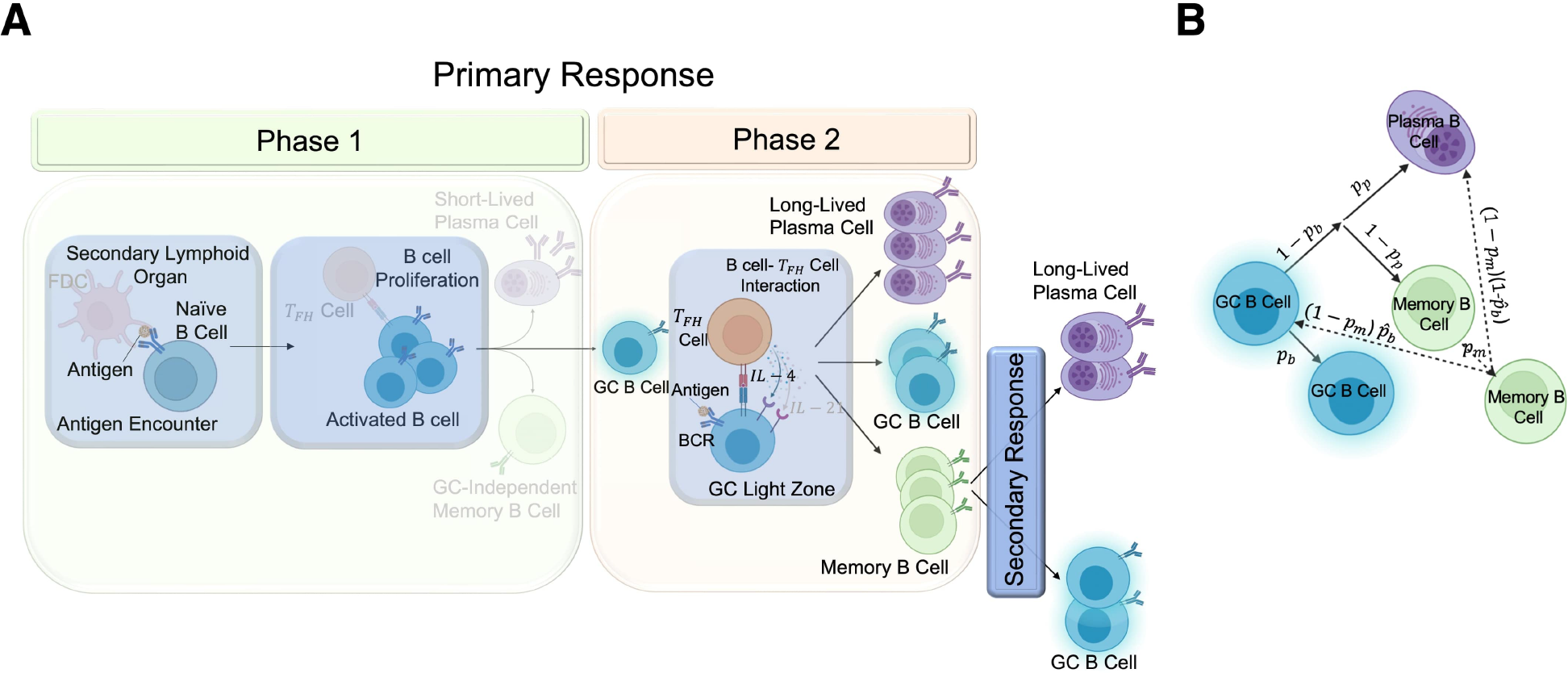
A) Development of the humoral immune response. In phase 1 of the primary response (left), after encountering antigen, signalling via the B cell receptor (BCR) in the secondary lymphoid organ initiates naïve B cell proliferation and differentiation into germinal centre GC B cells (highlighted compartments). In phase 2 of the primary response (right), newly differentiated GC B cells form GCs and present antigen to T follicular helper cells in the light zone. T helper cells activate B cells through IL-4 (highlighted cytokine) signalling. Upon exit from the GC, B cells terminally differentiate into plasma cells, memory B cells, or re-enter the GC dark zone. In the secondary response (bottom right), memory B cells respond to antigens by differentiating into long-lived plasma cells or GC B cells, restimulating antibody production. B) Fate of B cells after primary and secondary infections. Following primary infection, a GC B cell generates one GC B cell with a probability of *p*_*b*_, one plasma B cell with a probability of (1 −*p*_*b*_)*p*_*p*_ and one memory B cell with a probability of (1−*p*_*b*_)(1−*p*_*p*_). During secondary infection, a memory B cell can divide into another memory B cell with a probability of *p*_*m*_, a GC B cell with a probability of 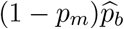, and a plasma B cell with a probability of 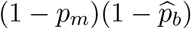.

#### Model Two

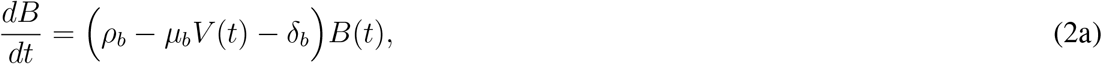

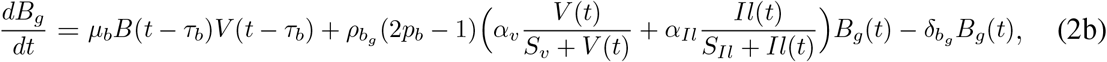

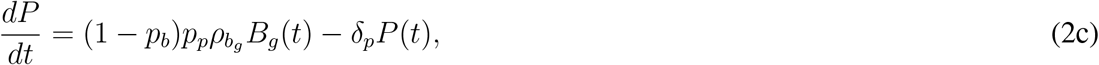

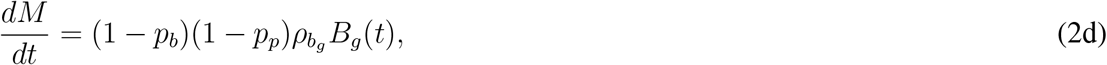

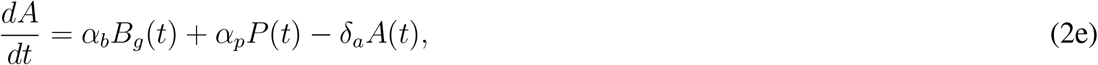

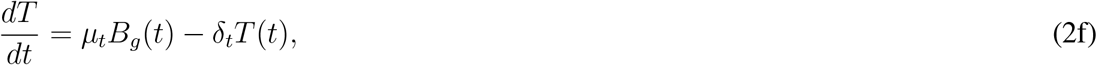

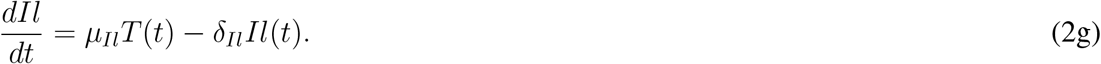

In Eq. (2a), *μ*_*b*_*V* (*t*) denotes the interaction of B cells with the pathogen (here considered to be SARS-CoV2 viral particles V(t)) and their subsequent differentiation at a rate of *μ*_*b*_, while *δ*_*b*_ represents the natural death rate of mature B cells.

Germinal centers are crucial in generating long-lived, high-affinity plasma and memory B cells [27, 28]. As the GC matures, B cells undergo multiple rounds of cell division, driven by interactions with T follicular helper cells and engagement with cognate antigens within the light zone [29]. In Eq. (2b), GC B cells are assumed to be activated after a delay of *τ*_*b*_ following naïve B cells first encounter with antigen. The activation of GC B cells depends on T follicular helper cells, IL-4 signals, and antigen interactions. The parameters *α*_*Il*_ and *S*_*Il*_ represent the binding rate of IL-4 to the receptor on the surface of GC B cells and the saturation constant of IL-4, respectively. The parameter *α*_*V*_ represents the binding rate of viral particles to GC B cell receptors, and *S*_*V*_ is the virus saturation constant. GC B cells undergo symmetric division with a rate of 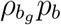 and die at a rate of 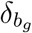.

Long-lived plasma B cells primarily form during the germinal center reaction and secrete high-affinity class antibodies [30, 31]. Eq. (2c) describes the differentiation of plasma B cells from GC B cells, which occurs with probability *p*_*p*_, and their natural death rate *δ*_*p*_. Memory B cells are long-lived and quiescent cells that respond upon re-stimulation by specific antigens [32–36]. They arise from the asymmetric division of GC B cells at a rate of 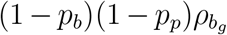 (Eq. (2d)). We assumed antibody production to be proportional to the number of GC B cells and plasma B cells and to occur at *α*_*b*_ and *α*_*p*_, respectively; antibodies degrade at rate *δ*_*a*_. This model focuses on neutralizing antibodies that can neutralize disease-causing pathogens, thereby providing immunity. T follicular helper cells play a crucial role in activating humoral immune responses. In our model, B cells act as antigen-presenting cells (APCs) for activating helper T cells in the light zone of germinal centers (Fig. 1). The term *μ*_*t*_*B*_*g*_(*t*) in Eq. (2f) represents the activation of *T* cells with a rate parameter *μ*_*t*_, based on the stimulation of GC B cells. The death of *T* cells is modelled by the term −*δ*_*t*_*T* with a death rate of *δ*_*t*_. Lastly, we consider IL-4 as the only cytokine in the system (Eq. (2g)). IL-4 is a cytokine with pleiotropic activity in the immune system [37], and it plays a crucial role in activating mature B cells. In our mathematical model, IL-4 is secreted by *T* cells at rate *μ*_*Il*_ and is cleared at rate *δ*_*Il*_. Table 2 summarizes model variables with their initial values. A detailed list of model parameter values and model variables with their respective initial values is given in Table 2.

**Table 2:**
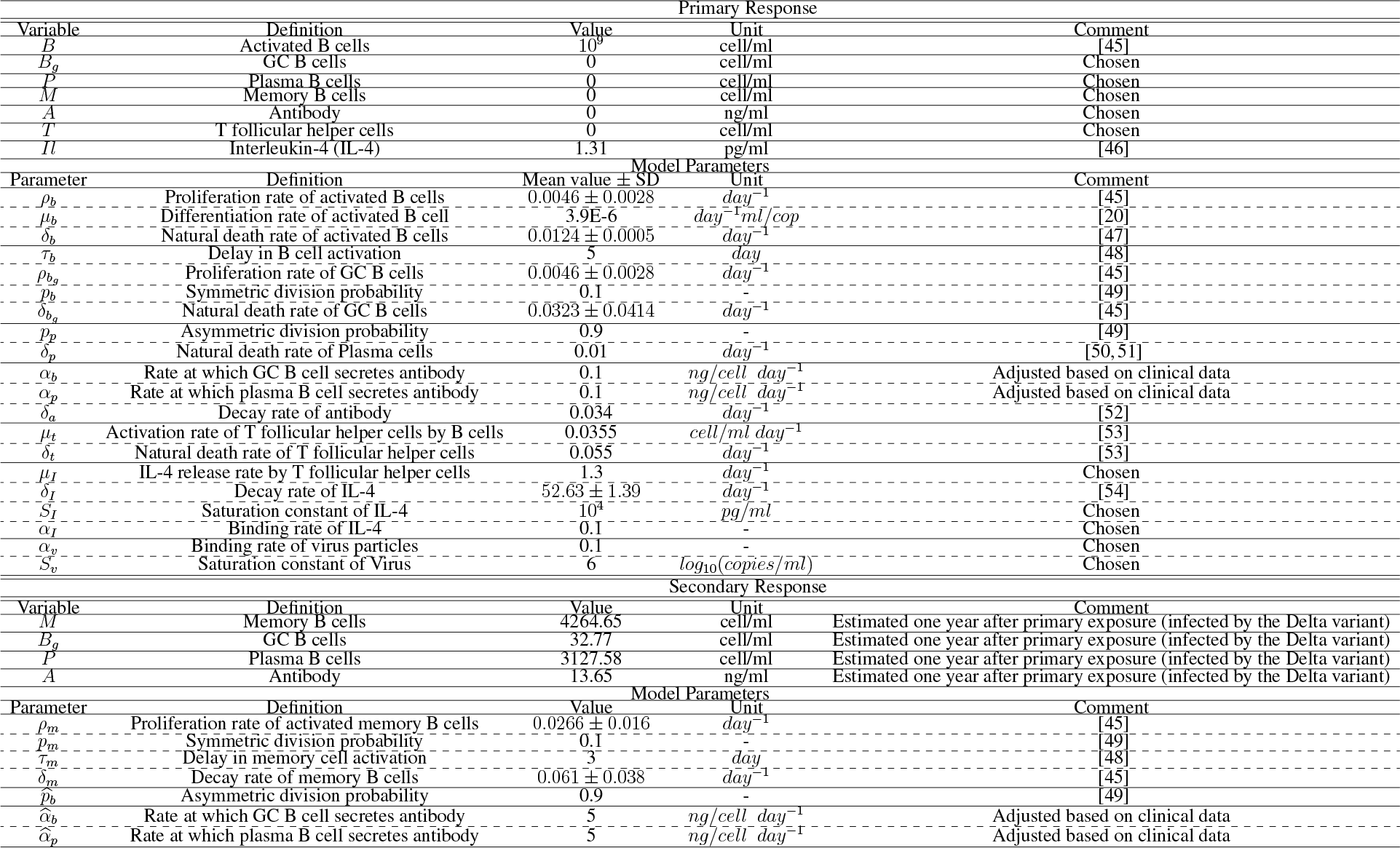
Model variables and parameters of primary and secondary adaptive immune responses (Eqs. (2) and (3)).

### Modeling re-exposure (secondary response)

Upon subsequent encounters with the same pathogen, the immune system can mount a faster and more robust response due to the previous establishment of immunological memory. This secondary immune response is typically effective in preventing disease by efficiently detecting, attacking, and eliminating the pathogen, leading to reduced symptoms. When memory B cells interact with their specific antigens upon re-exposure, they rapidly expand and generate a burst of plasma and germinal center B cells. To represent this evolving scenario, we formulated a mathematical model depicting the production of antibodies, as shown in Fig. 1. Within this model, we took into explicit account memory B cells, germinal center B cells, plasma B cells, and neutralizing antibodies. This model is characterized by a system of four nonlinear delay differential equations:

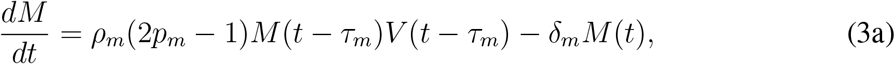

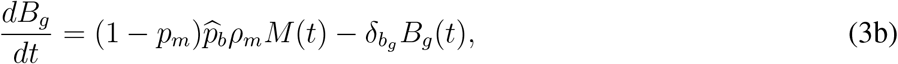

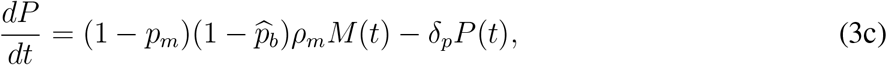

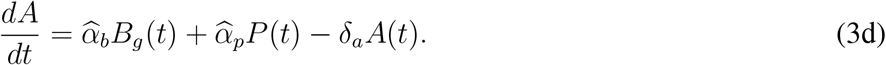

#### Model Three

In Eq. (3a), we allow for a short delay (*τ*_*m*_) to activate memory B cells after re-exposure to SARS-CoV-2 and consider mature memory B cells to die at rate *δ*_*m*_. Memory B cells undergo symmetric division at a rate *ρ*_*m*_*p*_*m*_. Eq. (3b) describes the dynamics of GC B cells, which are produced through differentiation of memory B cells with the probability of 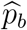. These GC B cells die naturally at rate 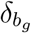. Long-lived plasma B cells, representing memory plasma B cells, are generated from the asymmetric division of memory B cells with a rate of 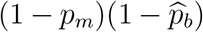, as shown in Eq. (3c).

The secondary antibody response is characterized by producing significant amounts of higher affinity IgG antibodies [38]. Therefore, we assumed different antibody generation rates for GC B cells 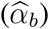 and plasma B cells 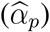 compared to the primary response (Eq. (3d)). The initial values of the secondary response depend on the specific day of re-exposure to the antigen, reflecting the time elapsed since the primary immune response. For example, the initial values for the secondary immune response at the one-year mark since the primary exposure (specifically, *M* (360), *B*_*g*_(360), *P* (360), *A*(360)), which were obtained by solving the primary response (Eqs. (2)) at day=360 (one year), are listed in Table 2.

### Antibody Neutralization Effect

Neutralizing antibodies play a critical role in the immune response by binding to specific regions (epitopes) on invading viruses, effectively neutralizing viral infections. They achieve this by blocking the interaction between the viral envelope and the host cell’s receptor or inhibiting the release of the viral genome [39]. To incorporate the impact of antibody neutralization in our model for SARS-CoV-2 infection within the host, we introduced an additional term in Eq. (1a) that accounts for the neutralizing effect of antibodies. This term enhances the inhibition of viral replication, reflecting the neutralization function. The modified equation is given by:

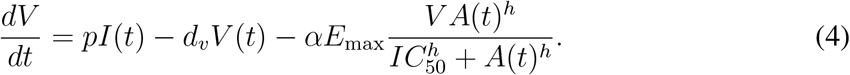

The parameters *E*_max_, *h*, and *IC*_50_ describe the neutralization function, which is crucial in determining the effectiveness of antibody neutralization and blocking new infections. *E*_max_ characterizes the maximal attainable neutralization achieved by antibodies and typically ranges between 0 and 1 (or 0 and 100%). Parameter *h* describes the gradient of the neutralization curve (usual Hill coefficient), signifying the degree of sensitivity in response to shifts in antibody concentration. *IC*_50_ represents the antibody concentration needed to achieve 50% neutralization.

Thus, by substituting Eq. (4) with Eq. (1a) in the within-host model (Eqs. (1)) and integrating it with either the primary humoral response model (Eqs. (2)) or the secondary humoral response model (Eqs. (3)), we can effectively simulate the interactions between the host and pathogen, and the stimulated immune response following the primary or secondary response, respectively. Notably, throughout this paper, we consistently utilized viral load dynamics that were influenced by the neutralization function (i.e., Eq. (4)).

### Model Calibration

#### Literature-Derived Parameters

Most of the parameters in our model were obtained from relevant literature sources. These fixed parameters represent constants that are well-established or values that have been empirically validated. A comprehensive list of the parameters used in our host-pathogen interaction model (as defined in Eqs (1)) is given in Table 1, with a reference to the source [23]. In the study by Jenner et al. (2021) [23], model parameters were obtained through various means, including direct extraction from existing literature, fitting of effect curves to experimental data collected in vitro, in vivo, and from clinical observations, or through the calculation of values that maintain homeostasis in the absence of SARS-CoV-2 infection. Furthermore, the parameters used to describe the immune response (as described in Eq. (2) and (3)) are meticulously detailed in Table 2, accompanied by the corresponding references.

Macallan et al. (2005) conducted a comprehensive study on the kinetics of human B lymphocytes, examining two distinct cohorts: one consisting of young individuals (below 35 years of age) and the other comprising elderly individuals (over 65 years of age), all in good health. Their observations revealed that peripheral blood B cells exhibited a relatively slow division rate, approximately 0.46% per day, while memory cells displayed a more rapid proliferation rate, approximately 2.66% per day (*ρ*_*m*_ = 0.0266). In the absence of specific data, we made the assumption that the proliferation rates for activated B cells and germinal center (GC) B cells were equivalent and set at 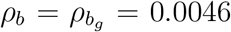. In the study by Perelson et al. (1976), biologically plausible parameter values were employed, assuming that B-lymphocytes were triggered and proliferated with a probability of 0.1. Consequently, in our model, we also assumed the same probability of symmetric division for activated B cells and activated memory B cells (*p*_*b*_ = *p*_*m*_ = 0.1).

#### Estimated Parameters

To determine the parameters associated with the neutralization (Equation (4)), namely {*E*_*max*_, *h, IC*_50_}, we used the curve-fit() function, a tool for nonlinear least squares curve fitting available within the Python programming language through the open-source SciPy library. The model was fit to data reporting the efficacy of clinical monoclonal antibodies (such as Sotrovimab) against the Delta and Omicron variants of SARS-CoV-2 from Planas et al. ([40]). We minimized the residual sum of squares (RSS):

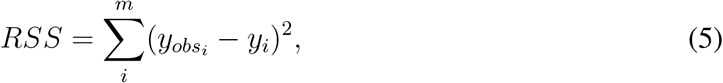

where the parameter *m* signifies the number of observed antibody concentration data points. We obtained the best-fit values for *E*_*max*_, *h*, and *IC*_50_ by minimizing the RSS between our model’s predictions and these data.

#### Adjusted parameters

Kinetic rates for the generation of antibodies from germinal center B cells and plasma cells are difficult to measure experimentally and are therefore generally unavailable. Thus, we leveraged data of the primary antibody response from eight hospitalized SARS-CoV-2 infected patients in Washington State, USA [41] to adjust the parameters *α*_*b*_ and *α*_*p*_ (primary response) in addition to 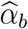 and 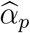 (secondary response) to ensure that model predictions captured the heterogeneity in antibody responses. For this, we simulated our model with parameters set as described in the previous sections and performed a parameter sweep ranging from 10^*−*4^ to 5. We then compared the model’s prediction to these data through a visual predictive check.

#### Sensitivity analysis

We conducted a global sensitivity analysis to identify the parameters most affecting antibody production to assess the impact of parameter variations on the maximum values of antibodies within both the primary and secondary immune responses in our mathematical model. We used Latin Hypercube Sampling (LHS) [42, 43] to generate 1000 samples of the model’s parameters. For each, we defined a parameter range using minimum = 0.5 × baseline parameter and maximum = 1.5 × baseline parameter. We used correlation and scatter plots to investigate the relationship between maximum antibody concentrations and parameters. Further, we measured the linear regression between predicted maximum antibody levels and changes in each parameter to elucidate the nature and strength of the relationship.

## Results

### Model calibration outcomes

#### Fitting neutralization function to clinical data

We performed curve fitting to clinical data from patients infected with the Omicron and Delta variants [40] to determine the parameters of the neutralization function (Eq. (4)). Separate curve fitting procedures were carried out for each variant, enabling us to extract variant-specific parameter values (Fig. 2A). The resulting parameter values are detailed in Table 1. The fitted parameters from the function suggest differences in the neutralization effect of antibodies against the Omicron and Delta variants of SARS-CoV-2. Notably, though we found a higher *E*_max_ value for Omicron (92.02) compared to Delta (83.94), indicating a higher maximum effect when antibody concentrations are at their saturating levels, the *IC*_50_ value for Omicron was found to be considerably higher (760.43) than for Delta (189.83), implying that a much greater concentration of antibodies is needed to achieve half of the maximum neutralization effect for Omicron. This suggests that Omicron is less susceptible to neutralization by the antibodies than Delta. The parameter *h*, or the Hill coefficient, further informs this interpretation. The Hill coefficient for Delta was estimated to be slightly above 1 (1.04), suggesting a cooperative binding. In contrast, for Omicron, the Hill coefficient was found to be less than 1 (0.84), which could indicate a negative cooperative effect or simply a lower level of cooperativity in antibody binding. Overall, the fitting results imply that while the maximum potential neutralization effect for Omicron may be higher, it is harder to achieve due to the need for higher antibody concentrations, indicating that Omicron may be more resistant to neutralization by antibodies than the Delta variant.

**Figure 2.**
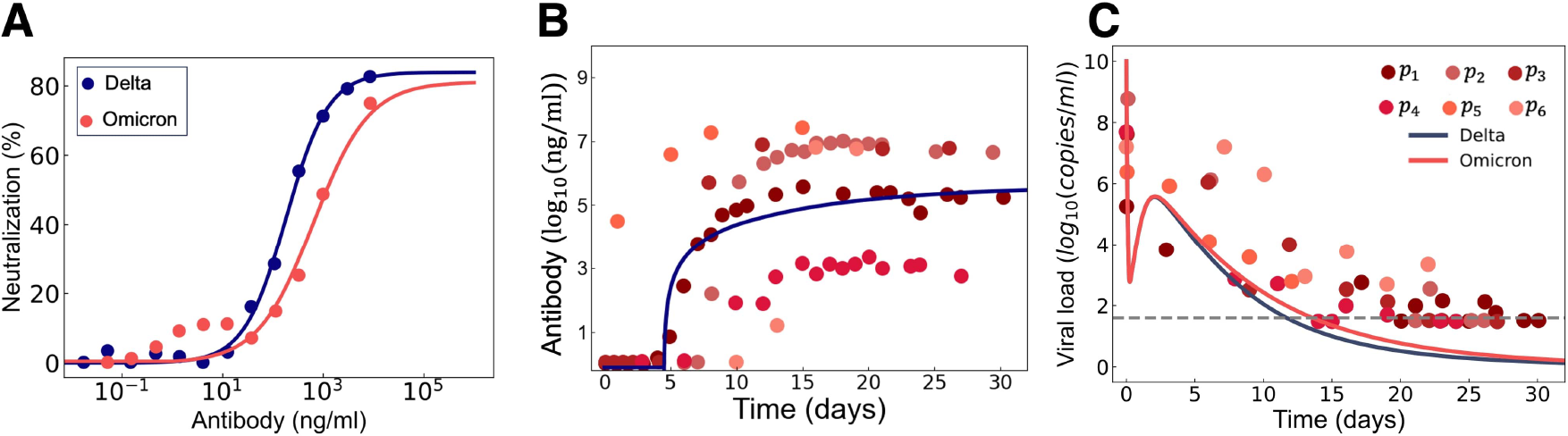
Comparisons of model predictions to clinical data. A) Model fits (solid lines) to the neutralization effect of monoclonal antibodies against SARS-CoV-2 Delta (blue dots) and Omicron (red dots) variants. B) Model prediction (blue solid line) of antibody concentrations in primary infections compared to clinical data from 8 hospitalized patients infected with SARS-CoV-2 Wuhan strain in Washington, USA (red markers) with *α*_*b*_ = 0.1 and *α*_*p*_ = 0.1. C) Model predictions of viral loads after infection by Delta (blue solid curve) or Omicron (red solid line) compared to the data. Horizontal dashed line: detection limit of 40 copies/ml.

#### Model validation

To validate the predictive capabilities of our model, we compared model predictions to clinical data collected from a cohort of eight hospitalized patients with SARS-CoV-2 infections [41]. This validation aimed to substantiate the accuracy of our model’s predictions pertaining to both antibodies in the primary response (Eq. (2e)) and viral load dynamics (i.e. Eq. (4)).

##### Antibody concentrations

The model’s predictions closely matched the measured antibody concentrations (Fig. 2B). To achieve this alignment, we set the values for the antibody generation rates of germinal center B cells and plasma B cells, represented by parameters *α*_*b*_ and *α*_*p*_, to 0.1. This choice allowed us to achieve a close correspondence between the model’s predictions and the clinical data. These data are from the first wave of the pandemic during which both variants had not yet emerged. For simplicity, we assumed Delta to be most similar to the Wuhan strain, given the evolutionary distance of Wuhan to Delta versus Wuhan to Omicron, and adjusted the parameters using the Delta prediction.

##### Viral load

Model predictions to data from SARS-CoV-2 concentrations from hospitalized patients encompassing infections from Delta and Omicron variants demonstrated good agreement. We found an elevation in viral loads associated with the Omicron variant compared to Delta (Fig. 2C).

### Antibody levels are strongly influenced by germinal centre and plasma B cell antibody generation rates

To quantify the influence of specific parameters on our predicted outcomes, we performed a global sensitivity analysis that focused on peak antibody concentrations (*A*_*max*_) after primary and secondary responses. The relationships between primary/secondary antibody responses and model parameters are depicted in Fig. 3A. In this figure, we have excluded parameters that do not have discernible impacts on model variations. During the primary immune response, our analyses reveal a weak correlation between the antibody generation rate by plasma B cells (*α*_*p*_) and the peak antibody concentration. In contrast, the antibody generation rate by GC B cells (*α*_*b*_) demonstrated a strong positive correlation. Furthermore, we observed a nearly equivalent negative correlation between the death rate of plasma B cells (*δ*_*p*_) and the clearance rate of anti-bodies (*δ*_*a*_) and the peak antibody value. In the secondary response, we found a similar positive correlation between the peak antibody value (*A*_*max*_) and the antibody generation rates of both GC B cells 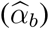 and plasma B cells 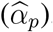. Moreover, *A*_max_ was strongly negatively correlated with the antibody clearance rate. This negative correlation was also evident, with a reduced coefficient value, between the death rates of memory B cells (*δ*_*m*_) and GC B cell level. Intriguingly, our findings also unveiled a positive correlation between *A*_*max*_ and the probability of symmetric deviation in memory B cells (*p*_*m*_). In contrast, a negative correlation was found with the probability of asymmetric deviation in memory B cells 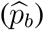. The observations from Fig. 3A were illustrated using scatter plots between the 1000 samples of each parameter generated through Latin hypercube sampling and the maximal predicted antibody concentration (Fig. 3B). Notably, a linear regression analysis yielded a higher Spearman’s correlation coefficient (*r* = 0.562) between *A*_*max*_ and *α*_*b*_ during the primary response, as compared to a correlation coefficient of *r* = 0.147 for *α*_*b*_. Similarly, in the secondary response, we found a strong correlation of *r* = 0.418 between *A*_*max*_ and 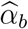, similar to the correlation *r* between *A*_*max*_ and 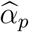 of *r* = 0.38. Low *p*-values (*<* 0.001) are reported in the regression fits.

**Figure 3.**
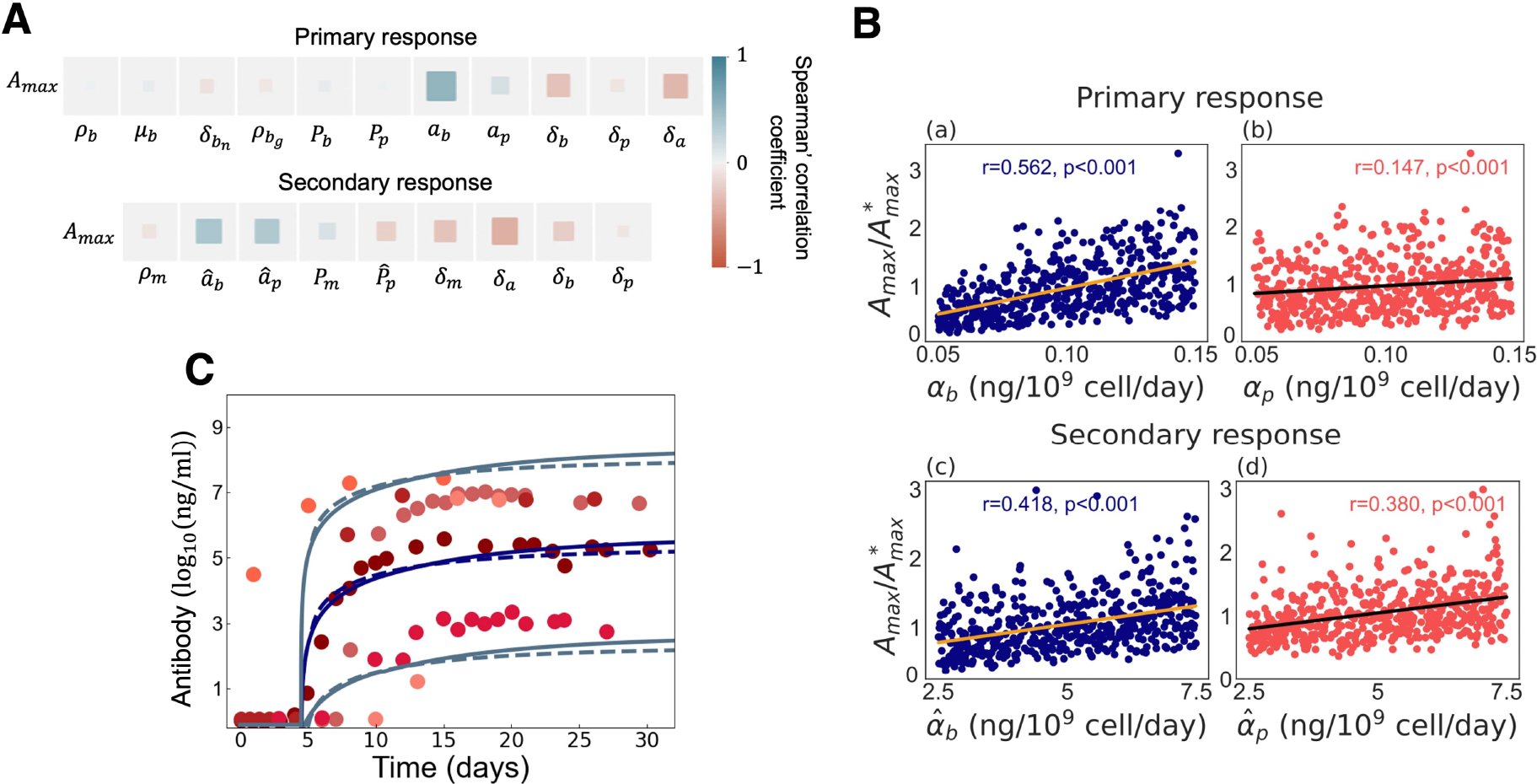
Identifying model parameters that significantly impact the maximum antibody level in both primary and secondary responses. A) Spearman’s rank correlation coefficient was calculated between the maximum primary and secondary antibody levels and model parameters. The blue and red colours indicate positive and negative correlations, respectively. The magnitude of the blue and red rectangles corresponds to the absolute value of the correlation rank, showing the statistical significance. B) Scatter plots with linear regression lines and Spearman’s correlation coefficients (*r* and *p*-value) are displayed for the primary and secondary antibody responses against (a, c) the antibody-secreting rate by germinal center B cells (*α*_*b*_ and 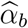) and (b, d) plasma B cells (*α*_*p*_ and 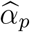). In (a) and (c), the golden lines represent the linear regression lines, while in (b) and (d), the black lines indicate the linear regression lines. The maximum antibody levels are normalized to the baseline values in primary and secondary responses. C) The model’s prediction of antibodies is compared with the clinical trial data from hospitalized patients. The gray lines in the graph depict the lowest and highest antibody levels captured by the model. The solid lines represent the primary response, while the dashed lines indicate the secondary response. The secondary exposure was modelled to occur one year after primary infection.

### Quantifying antibody generation rates in primary and secondary responses: model validation using clinical data

Given the above results from our sensitivity analysis, we next sought to capture the heterogeneity in antibody concentrations after primary infections reflected in the clinical data (see Fig. 2B). To recover the minimum and maximum values observed in the data from eight hospitalized COVID-19 patients [41], we modulated the baseline estimated values of parameters *α*_*b*_ and *α*_*p*_ in the primary response, along with 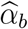 and 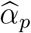 in the secondary response. Setting *α*_*b*_ = *α*_*p*_ = 10^*−*4^, our model predicted primary antibody concentrations from 10^2^ to 10^3^ (minimal observed values). Setting these two parameters to be *α*_*b*_ = *α*_*p*_ = 0.1 resulted in intermediate antibody levels ranging from 10^4^ to 10^6^. To achieve the highest antibody levels ranging from 10^7^ to 10^9^ required increasing both to *α*_*b*_ = *α*_*p*_ = 5 (Fig. 3C). Achieving the same antibody level following a secondary infection required a substantial increase in the parameters 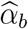 and 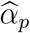, approximately 50 times higher than *α* and *α* (i.e., 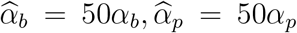). Notably, due to the lack of data detailing the distinctions between *α*_*b*_ and *α*_*p*_ (as well as 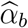 and 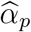 in the secondary immune response), we opted to set them to be equal. Furthermore, as described in subsection (), our dataset originates from eight hospitalized patients who had encountered primary SARS-CoV-2 infections; however, we also assessed the secondary antibody response.

### Antibody neutralization efficacy against Delta and Omicron variants

Using our full model with parameters values set to those in Table 2, we examined neutralization (Eq. (4)) in the context of the Delta and Omicron variants and found that neutralization (i.e., antibody efficacy) was higher during a secondary infection with Delta versus Omicron (Fig. 4A). This disparity implies that the Omicron variant may exhibit partial or complete neutralization evasion by the antibodies integrated into our model. Our findings align with the known immune-evasive properties of Omicron [55, 56] through reduced antibody binding to the Omicron spike domain, indicative of neutralization escape [40]. We also conducted a comparative analysis of antibody neutralization effects against Delta and Omicron secondary infections by considering three distinct scenarios for primary and secondary infections: (1) Delta-Delta, (2) Delta-Omicron, and (3) Omicron-Omicron infections, given a secondary infection occurring either three months, six months, or one year after the primary exposure to the virus (Fig. 4B). Our findings suggest that antibody neutralization is more pronounced in the Delta-Delta scenario compared to the other two scenarios, where similar neutralization effects were observed. Furthermore, the temporal interval between primary and secondary infections was found to have a strong influence on predicted neutralization, with delayed secondary infections resulting in reduced neutralization. This phenomenon can be attributed to the decreased antibody levels observed in Fig. 4C, coupled with an elevated viral load (Fig. 4D) during later-stage infections, and explains the increasing susceptibility to reinfection with time that has been observed from the beginning of the COVID-19 pandemic. Indeed, our results demonstrate higher antibody concentrations during secondary infections occurring three or six months after primary exposure, in contrast to secondary infections one year (Fig. 4C), indicative of waning immunity. Notably, we couldn’t detect any significant time differences in the decrease of neutralization effects for the various scenarios. In other words, there is no significant variation in the time intervals when neutralization reaches zero, whether for primary or secondary responses.

**Figure 4.**
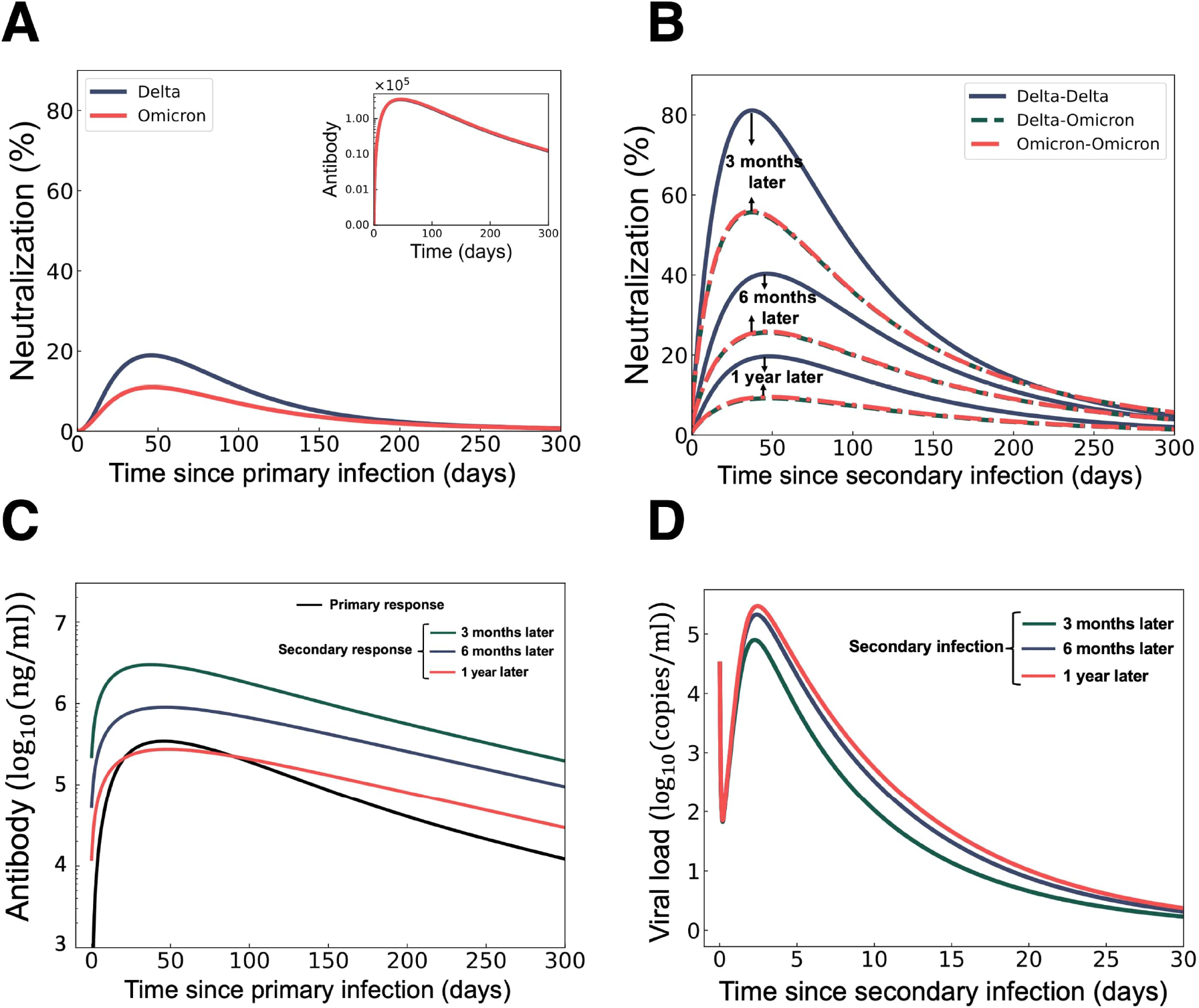
Predicting antibody neutralization effects on SARS-CoV-2 Delta and Omicron variants in primary and secondary immune responses. A) Primary response for Delta variant neutralization (blue curve) and Omicron variant neutralization (red curve). The inset reflects anti-body dynamics. B) Neutralization responses after 1) Delta-Delta infection (blue solid curve), 2) Delta-Omicron infection (dashed green curve), and Omicron-Omicron infection (dashed red curve), with secondary infections occurring three months, six months, and one year after the primary infection. C) Antibody responses in primary (black curve) and secondary infections taking place at three months (green curve), six months (blue curve), and one year (red curve) after the primary infection (Delta-Delta Scenario). D) Viral loads in Omicron secondary infection occurring three months (green curve), six months (blue curve), and one year (red curve) from the primary infection with Delta variant.

Our model simulations show that waning waning antibody levels can be attributed to the decreasing populations of germinal center and plasma B cells. This trend is evident in Fig. 5, where the initial quantities of GC B cells (Fig. 5A) and plasma B cells (Fig. 5B) are considerably smaller and continue to decrease over time. In other words, their initial values one year after the primary infection are smaller than those at six months and significantly lower than the levels observed at three months (e.g. (*B*_*g*_(90) *> B*_*g*_(180) *> B*_*g*_(360)) and (*P* (90) *> P* (180) *> P* (360)).

**Figure 5.**
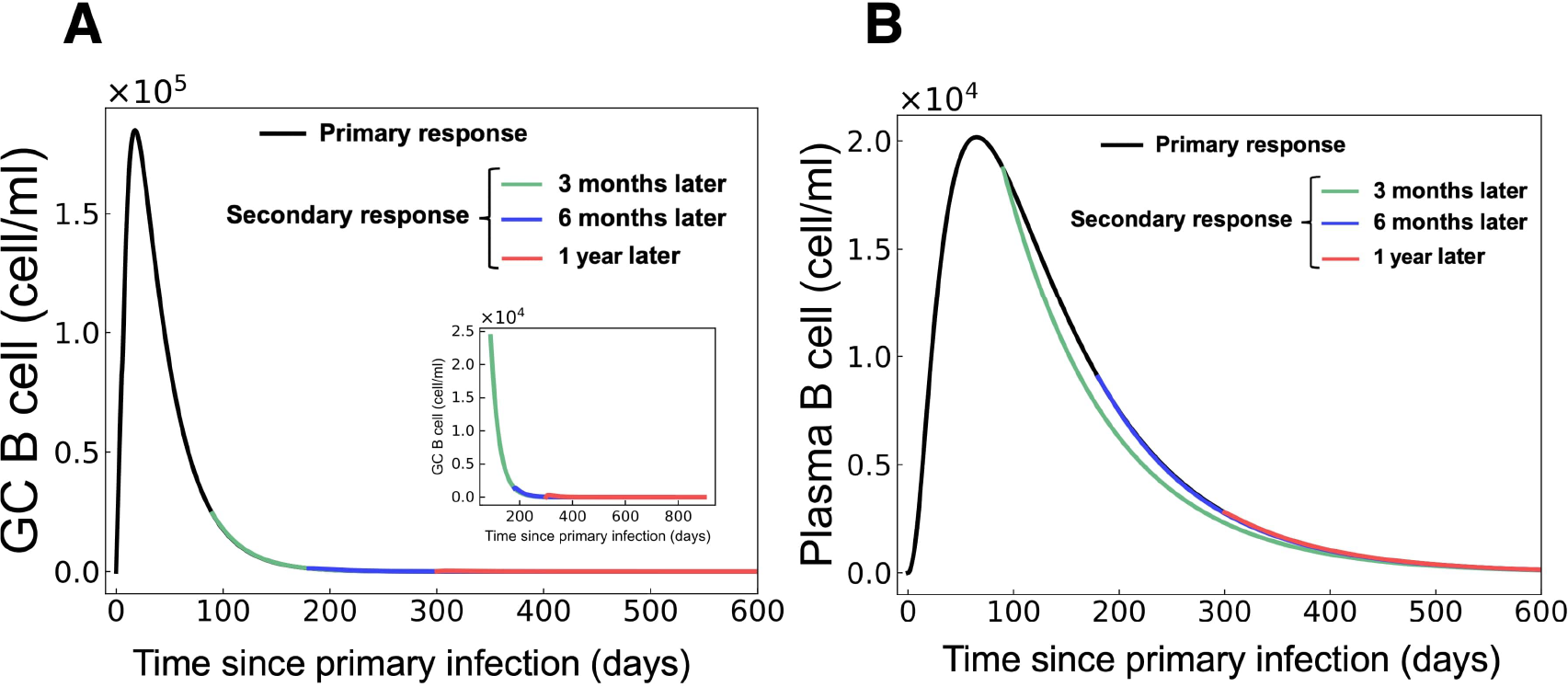
Immune cell dynamics post primary and secondary infections with Delta variant for A) GC B cells and B) plasma B cells over time. Black: primary response. Green: secondary response to infection occurring three months after primary. Blue: secondary response to infection occurring six months after the primary. Red: secondary response to infection occurring one year after primary. The inset reflects GC B cell dynamics in the secondary immune response.

## Discussion

In this study, we developed a novel mathematical model to explore the intricate processes governing B lymphocyte activation, replication, and differentiation and the production of antibodies during infection by SARS-CoV-2. Our model traces the path from germinal center B cells to memory and long-lived plasma B cells, culminating in the production of antibodies after initial and subsequent viral exposures using a system of delay differential equations (DDEs) to capture the interactions between immune cells and neutralizing antibodies. We assumed a delay in the activation of germinal center B cells during the primary immune response and a shorter delay in the activation of memory B cells during the secondary immune response. By incorporating the concept of neutralization, characterized by the binding of antibodies to viral particles to hinder their replication, we could study the antiviral potency exhibited by neutralizing antibodies. Neutralizing antibodies are crucial in mitigating viral infectivity and have been widely studied as potential therapeutic agents [57]. Although a substantial portion of our model parameters were sourced from existing literature, specific parameters required informed assumptions. For this, we conducted a comprehensive global sensitivity analysis to unveil the parameters exerting significant influence over the outcomes of our model. Our research revealed that in the primary response, the maximum antibody level was most sensitive to the antibody generation rate by germinal center B cells. In contrast, it was most sensitive to antibody production rates of both germinal center B cells and plasma B cells in the secondary response. By modulating these parameters and comparing model predictions to clinical data, we found that higher antibody generation rates in the secondary immune response are needed to reach comparative antibody concentrations in both primary and secondary infections.

Investigating the neutralizing effect of antibodies against Delta and Omicron variants after primary infection revealed a diminished neutralization rate for Omicron despite parity in antibody levels. This finding corresponds to known immune evasive properties of the Omicron variant [58–60]. We noted a declining trend in overall neutralization when evaluating the secondary immune response at intervals of three, six, or one year following primary infection. This trend aligns with the decrease in antibody levels, which is a consequence of the reduced initial GC and plasma B cells, predicted by our model, thus indicating its capacity to capture waning immunity. Waning immunity has particular importance for vaccination campaign scheduling. Therefore, beyond the essential biological insights gained from this work, our model could be used in public health contexts for planning boosters.

In summary, our model of the humoral response predicted (1) antibody and viral load dynamics for various SARS-CoV-2 variants, such as Delta and Omicron, in agreement with clinical patient data, (2) elevated secondary immune responses characterized by augmented antibody generation rates by germinal center B cells and plasma B cells, coupled with intensified antibody neutralization effects, (3) the immune-evasive nature of the Omicron variant, marked by similar antibody levels but higher viral load and diminished neutralization tendencies compared to the Delta variant and (4) waning immunity. This study thus contributed to our understanding of humoral immunity to SARS-CoV-2 and other respiratory viruses and can be used to predict antibody dynamics following infection or vaccination. It is important to note that our immune models do not take into account affinity coefficients for antibodies. Therefore, we relied on variations in the antibody generation rate by germinal center B cells and plasma B cells to capture differences in antibody concentrations between the primary and secondary responses. This limitation highlights the need for further refinement and expansion of our model to incorporate additional factors, such as affinity maturation. While our study primarily focused on B cells, antibodies, and the contributions of memory B cells to the secondary response, the adaptive immune response is a complex interplay of various molecules and cell types, including T cell-mediated immunity. Future studies will explore these factors and refine our modelling approach accordingly. Lastly, it should be noted that since we used a deterministic framework consisting of ordinary and delay differential equations, our model will predict viral titers below the threshold of a cleared infection (generally considered to be 1 − 2 *log*_10_(*copies/ml*)), unlike stochastic systems. Overall, our mathematical model provides valuable insights into the dynamics of humoral immunity and the role of neutralizing antibodies in the context of SARS-CoV-2 infection. By uncovering our model’s critical parameter values and limitations, we lay the foundation for future investigations to understand better the adaptive immune response following SARS-CoV-2 infections and reinfections with matched or discordant strains and potential therapeutic interventions.

## Acknowledgments

The Authors acknowledge support from the One Health Modelling Network for Emerging Infections, funded by the Natural Sciences and Engineering Research Council of Canada (NSERC) and the Public Health Agency of Canada (PHAC) as part of the Emerging Infectious Disease Modelling Initiative, NSERC Discovery Grant RGPIN-2018-04546 (MC), the Fondation du CHU Sainte-Justine (MC), and a Fonds de recherche du Québec-Santé J1 Research Scholar Award (MC).

